# Tractography-based connectomes are dominated by false-positive connections

**DOI:** 10.1101/084137

**Authors:** Klaus H. Maier-Hein, Peter Neher, Jean-Christophe Houde, Marc-Alexandre Côté, Eleftherios Garyfallidis, Jidan Zhong, Maxime Chamberland, Fang-Cheng Yeh, Ying-Chia Lin, Qing Ji, Wilburn E. Reddick, John O. Glass, David Qixiang Chen, Yuanjing Feng, Chengfeng Gao, Ye Wu, Jieyan Ma, H Renjie, Qiang Li, Carl-Fredrik Westin, Samuel Deslauriers-Gauthier, J. Omar Ocegueda González, Michael Paquette, Samuel St-Jean, Gabriel Girard, François Rheault, Jasmeen Sidhu, Chantal M.W. Tax, Fenghua Guo, Hamed Y. Mesri, Szabolcs Dávid, Martijn Froeling, Anneriet M. Heemskerk, Alexander Leemans, Arnaud Boré, Basile Pinsard, Christophe Bedetti, Matthieu Desrosiers, Simona Brambati, Julien Doyon, Alessia Sarica, Roberta Vasta, Antonio Cerasa, Aldo Quattrone, Jason Yeatman, Ali R. Khan, Wes Hodges, Simon Alexander, David Romascano, Muhamed Barakovic, Anna Auría, Oscar Esteban, Alia Lemkaddem, Jean-Philippe Thiran, H. Ertan Cetingul, Benjamin L. Odry, Boris Mailhe, Mariappan S. Nadar, Fabrizio Pizzagalli, Gautam Prasad, Julio E. Villalon-Reina, Justin Galvis, Paul M. Thompson, Francisco De Santiago Requejo, Pedro Luque Laguna, Luis Miguel Lacerda, Rachel Barrett, Flavio Dell’Acqua, Marco Catani, Laurent Petit, Emmanuel Caruyer, Alessandro Daducci, Tim B. Dyrby, Tim Holland-Letz, Claus C. Hilgetag, Bram Stieltjes, Maxime Descoteaux

**Author notes:** indicates corresponding authors.

## Abstract

Fiber tractography based on non-invasive diffusion imaging is at the heart of connectivity studies of the human brain. To date, the approach has not been systematically validated in ground truth studies. Based on a simulated human brain dataset with ground truth white matter tracts, we organized an open international tractography challenge, which resulted in 96 distinct submissions from 20 research groups. While most state-of-the-art algorithms reconstructed 90% of ground truth bundles to at least some extent, on average they produced four times more invalid than valid bundles. About half of the invalid bundles occurred systematically in the majority of submissions. Our results demonstrate fundamental ambiguities inherent to tract reconstruction methods based on diffusion orientation information, with critical consequences for the approach of diffusion tractography in particular and human connectivity studies in general.

---

Fiber tractography, a computational reconstruction method based on diffusion-weighted magnetic resonance imaging (DWI), attempts to reveal the white matter pathways of the human brain *in vivo* and to infer the underlying network structure, the structural connectome (1). Numerous algorithms for tractography have been developed and applied to connectome research in the fields of neuroscience (2,3) and psychiatry (4). Given the broad interest in this approach, the advantages and shortcomings of tractography have been widely debated (1,5–9). Particularly, *in vivo* tractography of the human brain was evaluated by subjective assessment of plausibility (10) or qualitative visual agreement with *post-mortem* Klingler-like dissections (11,12). Reproducibility (13) or data prediction errors (14–16) were evaluated in the context of tractographic model verification. However, these evaluations cannot validate the accuracy of reconstructions (17). Moreover, *ex vivo* imaging and tracing (17–25) or physically (26–32) and numerically simulated phantoms (33–37) allow validation to some extent. The complexity of the nervous system, however, and the lack of precise ground truth information on the trajectories of pathways and their origins as well as terminations in the human brain make it hard to quantitatively evaluate tractography results and to determine which discoveries are reliable when regarding brain connectivity in health and disease.

State-of-the-art tractography algorithms are driven by *local orientation fields* estimated from DWI, representing the local tangent direction to the white matter tract of interest (1). Conceptually, the principle of inferring connectivity from local orientation fields can lead to problems as soon as pathways overlap, cross, branch and have complex geometries (Figure 1) (8,38,39). Since the invention of diffusion tractography, these problems have been discussed in schematic representations or theoretical arguments (8,9,40), but have not yet been quantified in brain imaging. To determine the current state of the art in tractography, we organized an international tractography competition (tractometer.org/ismrm_2015_challenge) and employed a novel validation method based on simulated DWI of a brain-like geometry. This ground truth data set represented 25 well-known valid bundles that covered approximately 70% of the human brain white matter.

**Figure 1.**
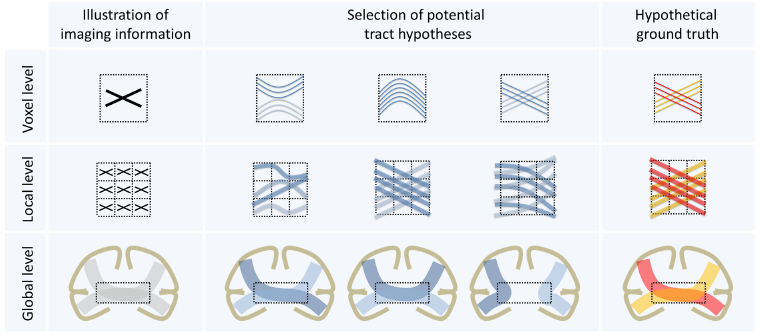
Ambiguous correspondences between diffusion directions and fiber geometry. The three illustrations at voxel, local and global level exemplarily illustrate ambiguities in apparent diffusion imaging information, leading to several potential tract reconstructions. The intra-voxel crossing of fibers in the hypothetical ground truth (first row) leads to ambiguous imaging information at voxel level **(8)**. Similarly, the imaging representation of local fiber crossings (second row) can be explained by several other configurations **(8)**. At a global level (third row), white matter regions that are shared by multiple bundles (“bottlenecks”, dotted rectangle) **(38)** can lead to many spurious tractographic reconstructions **(39)**. With only two bundles in the hypothetical ground truth (red and yellow bundle), four potential false-positive bundles emerge.

At the closing of the competition, we evaluated 96 distinct tractography pipelines submitted by 20 different research groups, in order to assess how well the algorithms were able to recover the known connectivity. We also assessed essential processing steps to pinpoint critical flaws that many current pipelines have in common. On average, submissions recovered only a third of the volumetric extent of existing bundles. Also, most algorithms routinely extracted many false positive bundles, even though they were not part of the ground truth. Some of these false-positive bundles resemble previously reported pathways identified by *in vivo* tractography, such as the frontal aslant tract (41) or the vertical occipital fasciculus (42). The average ratio of false-positive to true-positive bundles was approximately four to one. This ratio could not be improved by employing higher quality data or even using the gold standard field of local orientations, highlighting that current tractography approaches are fundamentally ill-posed.

## Results

### Datasets and submissions

Prior investigations of tractography methodology have chosen artificial fiber geometries to construct synthetic ground truth models (28,43). Here, we defined our ground truth based on the fiber bundle geometry of a high-quality Human Connectome Project (HCP) dataset that was constructed from multiple whole-brain global tractography maps (44) (Supplementary Figure 1). Following the concepts introduced in (45), an expert radiologist (B.S.) extracted 25 major tracts (i.e., bundles of streamlines) from the tractogram. These association, projection and commissural fibers covered more than 70% of the white matter across the whole brain. The dataset features a brain-like macro structure of long-range connections, mimicking *in vivo* DWI clinical-like acquisitions based on a simulated diffusion signal. An additional anatomical image with T1-like contrast was simulated as a reference. The final datasets and all files needed to perform the simulation are available online (Supplementary Data 1).

20 research groups across 12 countries (Supplementary Figure 2a) participated in the competition and submitted a total of 96 processing results comprising a large variety of tractography pipelines with different pre-processing, local reconstruction, tractography and post-processing algorithms (Supplementary Figure 2b, Supplementary Notes 2).

### Performance metrics and evaluation

The Tractometer connectivity metrics (43) were used for evaluating submissions. Based on the known ground truth bundles, we calculated true positives, corresponding to the valid connection ratio (VC), that is, the proportion of streamlines connecting valid end points and the associated number of valid bundles (VB), where a bundle is a group of streamlines. We also computed false positives, corresponding to the invalid connection ratio (IC) and the associated number of invalid bundles (IB), as well as reconstructed volumes, based on the bundle volumetric overlap (OL) and volumetric overreach (OR) in percent (Supplementary Figure 3).

### Tractography detects major bundles, but not to the full extent

While the tractography algorithms detected most existing bundles, the extent of their volumetric reconstruction varied. Figure 2a groups identified valid bundles into three clusters of “very hard”, “hard” and “medium” bundles. Figure 2b shows corresponding examples that were reconstructed by different tractography techniques. Most submissions had difficulties identifying the smallest tracts, that is, the anterior (CA) and posterior commissures (CP) that have a cross-sectional diameter of no more than two millimeters, or one or two voxels (“very hard”). A family of “hard” bundles was recovered by almost all algorithms, with a low overlap score of approximately 30%. Bundles of "medium" difficulty were the corpus callosum (CC), inferior longitudinal fasciculus (ILF), superior longitudinal fasciculus (SLF) and uncinate fasciculus (UF). These tracts span different shapes, lengths and sizes, but were recovered most frequently, with above 45% overlap and around 40% overreach.

**Figure 2.**
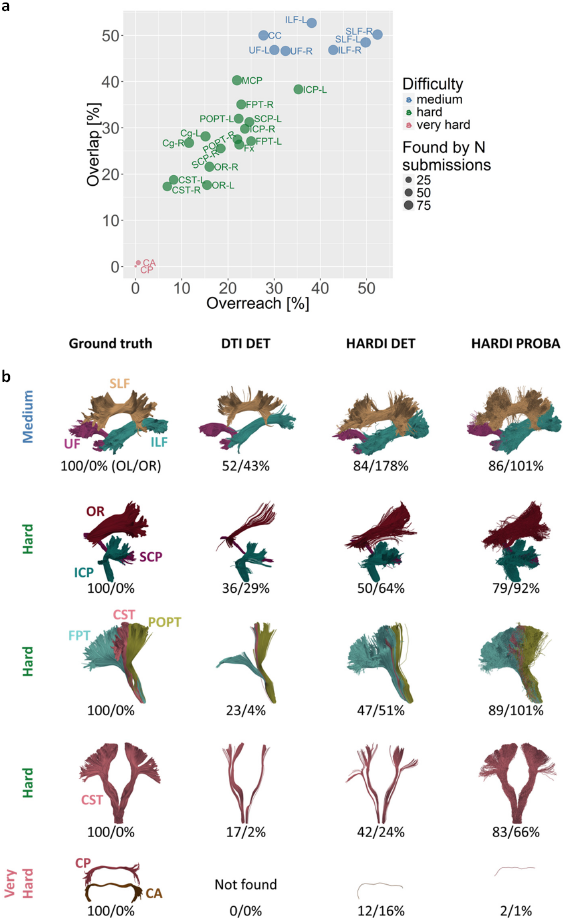
Tractography detects major bundles, but not to the full extent. (a) Overview of scores reached for different bundles in ground truth. Average overlap (OL) and overreach (OR) scores for the submissions (red: “very hard”, green: “hard”, blue: “medium”, for abbreviations see Supplementary Figure 1). (b) Representative bundles for diffusion tensor imaging (DTI) deterministic (DET) tracking come from submission 6 / team 20, high angular resolution diffusion imaging (HARDI) deterministic tracking from submission 0 / team 9 and HARDI probabilistic (PROBA) tracking from submission 2 / team 12 (see Supplementary Notes 5 for a discussion of these submissions). The first column shows ground truth valid bundles for reference. The reported OL and OR scores correspond to the highest OL score reached within the respective class of algorithms.

As shown in Figure 3, the submissions identified an average of 21 out of 25 valid bundles (median 23). No team detected all valid bundles, but three teams (10 submissions) recovered 24 valid bundles, and 15 out of 20 teams (69 submissions) detected 23 or more valid bundles (Figure 4a, red entries in connectivity matrix). However, tractography pipelines clearly need to improve their recovery of the full spatial extent of bundles (Figure 3c). The mean value of bundle volume overlap (OL) across all submissions was 31% ± 17%, with an average overreach (OR) of 23% ± 21%. At the level of individual streamlines, an average of 54% ± 23% connections were valid (Figure 3a).

**Figure 3.**
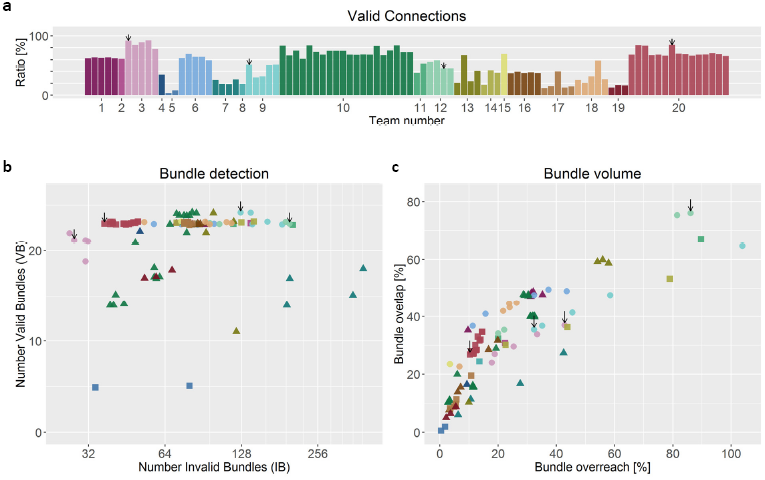
Tractography identifies more invalid than valid bundles. Overview of scores reached by the different teams. (a) Percentage of streamlines connecting valid regions. (b) Number of detected valid and invalid bundles (data points are jittered to improve legibility). (c) Volume overlap (OL) and overreach (OR) scores averaged over bundles. Black arrows mark submissions used in the following figures (see Supplementary Notes 5 for discussion).

**Figure 4.**
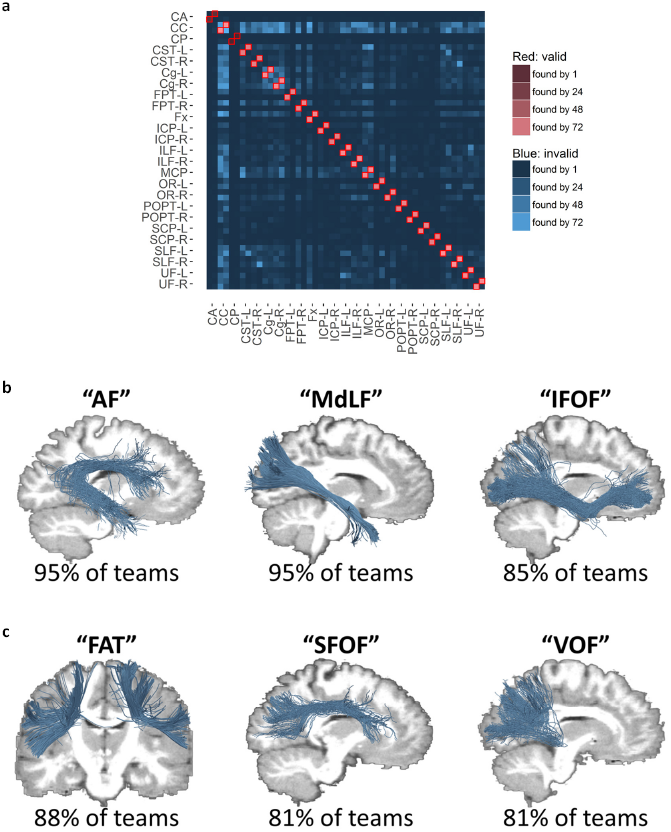
Overview of valid (red) and invalid (blue) bundles. Clusters of invalid streamlines exhibit similarities to previously reported, yet unconfirmed bundles from the literature. (a) Each entry in the connectivity matrix indicates the number of submissions that have identified the respective bundle. The two rows/columns per bundle represent the head-endpoint and tail-endpoint regions of a bundle. (b, c) The shown invalid bundles have been consistently identified by more than 80% of the teams, but do not exist in the ground truth. As discussed in the main text, some of these bundles exhibit a high similarity to bundles that are known to exist, as shown in (b), including the arcuate fasciculus (AF) and bundles traversing the temporal stem such as the middle longitudinal fasciculus (MdLF) and the inferior-frontal occipital fasciculus (IFOF). Other bundles – shown in (c) – are not known to exist and are debated in literature. These include the frontal aslant tract (FAT), the superior fronto-occipital fasciculus (SFOF) and the vertical occipital fasciculus (VOF).

### Tractography identifies more invalid than valid bundles

Across submissions, 36% ± 17% of the reconstructed individual streamlines connected regions that were not actually connected. The fraction of streamlines not connecting any endpoints was 10% ± 15%. Even though not part of the ground truth, these streamlines often occur in dense, structured and coherent bundles. On average, submissions identified more than four times as many invalid bundles as they identified valid bundles (Figure 3b). Submissions with at least 23 valid bundles showed no fewer than 37 invalid bundles (mean 88 ± 39, n = 69). Submissions with 23 or more valid bundles *and* a volumetric bundle overlap of > 50% identified between 99 and 204 invalid bundles (corresponding to more than four invalid bundles per valid bundle). The submissions produced an average of 88 ± 58 invalid bundles, demonstrating the inability of current state-of-the-art tractography algorithms to control for false positives (Figure 4a, blue entries in connectivity matrix). 41 of these invalid bundles occurred in the majority of submissions (Supplementary Figure 4).

The bundles illustrated in Figures 4b and 4c were systematically found by more than 80% of submissions without being part of the ground truth. Interestingly, several of these invalid streamline clusters exhibited similarities to bundles known or previously debated in tractography literature. They anatomically resemble the following pathways: 1) the frontal aslant tract (FAT) (41), 2) the arcuate fasciculus (AF) (46,47), 3) bundles passing through the temporal stem (48), such as the inferior-frontal occipital fasciculus (IFOF) (49,50), middle longitudinal fasciculus (MdLF) (51,52) and the extreme capsule fasciculus (EmC) (53), 4) the superior fronto-occipital fasciculus (SFOF) (49,54) and 5) the vertical occipital fasciculus (VOF) (42). The existence of the FAT, SFOF and VOF is controversial (41,42,49,54).

### Higher image quality does not solve the problem

To confirm that our findings revealed fundamental properties of tractography itself and are not related to effects of our specific phantom simulation process, we ran additional tractography experiments directly on the *ground truth field of fiber orientations* (Supplementary Figure 5 and Supplementary Notes 3), that is, without using the diffusion-weighted data at all. This setup was, thus, independent of image quality, artifacts and many other influences from specific pipeline configurations and the phantom generation process. Based on the ground truth orientations, the tractography pipelines achieved much higher overlap (71% ± 2%) and lower overreach (20% ± 0.2%) scores, while achieving valid connection ratios between 75% and 81%. However, they still generated more than three times as many invalid bundles than valid bundles (at least 82 invalid bundles).

### Tractography is fundamentally ill-posed

Our results show that the geometry of many junctions in the brain is too complex to be resolved by current tractography algorithms, even when given a perfect ground truth field of orientations. In the temporal lobe, for example, multiple bundles overlap and clearly outnumber the count of fiber orientations in most of the voxels. As illustrated in Figure 5, *single* fiber directions in the diffusion signal regularly represent multiple bundles (see also Supplementary Video 1). Such funnels embody hard bottlenecks for tractography, leading to massive combinatorial possibilities of plausible configurations for connecting the associated bundle endpoints as sketched in Figure 5c. Consequently, for the real dataset as well as the synthetic phantom, dozens of structured and coherent bundles pass through this bottleneck, exhibiting a wide range of anatomically reasonable geometries as illustrated in Supplementary Video 2. A tractogram based on real HCP data exhibits whole families of theoretically plausible bundles going through the temporal lobe bottleneck even though, locally, the diffusion signal often shows only one fiber direction (cf. Figure 5d).

**Figure 5.**
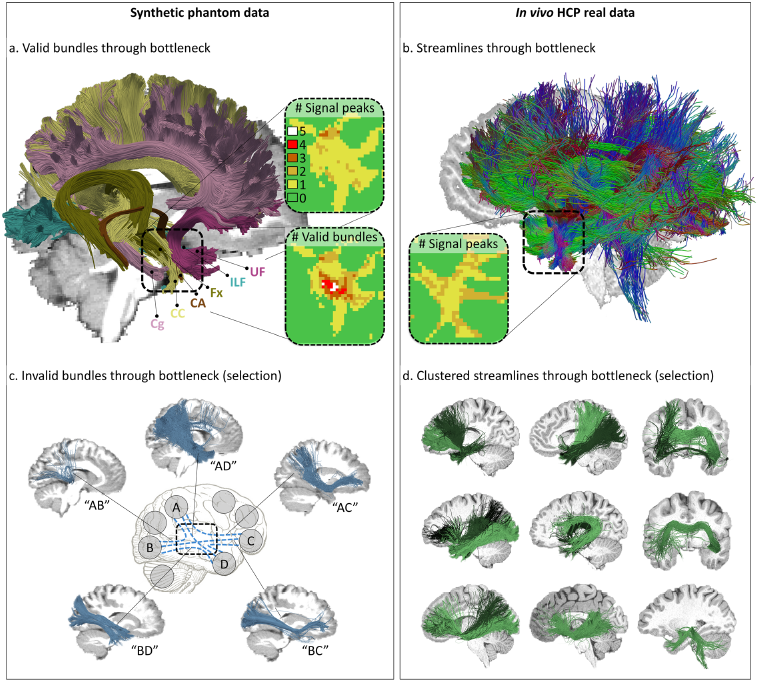
Fundamental ill-posedness of tractography. (a) Visualization of six ground truth bundles converging into a nearly parallel funnel in the bottleneck region of the left temporal lobe (indicated by square region). The bundles per voxel (box “# Valid bundles”) clearly outnumber the peak directions in the diffusion signal (box “# Signal peaks”). (b) Visualization of streamlines from a Human Connectome Project (HCP) *in vivo* tractogram passing through the same region. (c) Exemplary invalid bundles that have been identified by more than 50% of the submissions, showing that tractography cannot differentiate between the massive amount of plausible combinatorial possibilities connecting different endpoint regions (see Supplementary Video 1). (d) Automatically Quickbundle-clustered streamlines from the *in vivo* tractogram going through the temporal region of interest. The clustered bundles are illustrated in different shades of green. These clusters represent a mixture of true-positive and false-positive bundles going through that bottleneck area of the HCP data set (see Supplementary Video 2).

### Statistical analysis of processing steps

Effects of the methodological setup of the different submissions on the results were investigated in a multivariable linear mixed model and revealed the influence of the individual processing steps on the tractography outcome (Supplementary Table 1, Supplementary Notes 4). The choice of tractography algorithm, as well as the post-tracking filtering strategy and the underlying diffusion modeling had a strong effect on overall scores, revealing a clear tradeoff between sensitivity and specificity.

## Discussion

We assessed current state-of-the-art fiber tractography approaches using a ground truth dataset of white matter tracts and connectivity that is representative of the challenges that occur in human brain imaging *in vivo*. Advanced tractography strategies in combination with current diffusion modeling techniques successfully recovered most valid bundles, covering up to 76% of their volumetric extent. This sensitivity comes at a high cost. Tractography systematically also produced thick and dense bundles of plausible looking streamlines in locations where such streamlines did not actually exist. When focusing on the 64 bundles that were systematically recovered by the majority of submissions, 64% of them were in fact absent from the ground truth. Currently even the best tractography pipelines, or *tracking of the ground truth fiber orientations*, produce more falsepositive than true-positive bundles. These findings expose the degree of ambiguity associated with whole-brain tractography and show how the computational problem of tractography goes far beyond the local reconstruction of fiber directions (1,8) and issues of data quality. Our findings, therefore, present a core challenge for the field of tractography and connectivity mapping.

High invalid-connection ratios were previously reported under simplified conditions (28,43) (www.tractometer.org), and some of the underlying ambiguities in tractography have been discussed using schematic representations and theoretical arguments (1,8,9,40). Regions of white matter bottlenecks have been discussed in the past (38) and have been highlighted as critical with respect to tractographic findings (39). The present results reveal and quantify the consequences of such limitations under conditions found in human brain studies *in vivo*, addressing important questions that previously remained speculative. The findings were derived from a brain-like geometry that encompasses some of the major known long-range connections and covers more than 70% of the white matter. Future versions of the phantom are planned to include additional bundles such as the middle and inferior temporal projections of the AF, the MdLF and the IFOF as well as smaller U-fibers, medial forebrain fibers, deep nuclei and connections between them. In addition, more advanced diffusion modeling methods will allow generating even more realistic DWI signals, potentially simulated at increased spatial and q-space resolutions. These developments, however, will not resolve the fundamental ambiguities which tractography faces and thus will only have a limited effect on the main findings of our study. We showed that false-positive bundles occur at similar rates even when using the maximal angular precision of the signal, that is, using ground truth orientations. Increasing the anatomic complexity of the phantom by adding more bundles will most likely lead to even increased false-positive rates. The construction process of the current phantom resembles a potential limitation, since it involves tractography itself and thus raises self-validation issues. This should be considered in direct method comparisons as there may exist a possible bias towards algorithms that equal the algorithm used for phantom generation. This caveat, however, has only a very limited effect on our general findings. It can be expected that the identified limitations of tractography will become even more pronounced in phantoms of higher anatomic complexity that might be achievable by involving independent methods such as polarized light imaging (PLI) (55). In summary, our observations confirm the fundamental ill-posedness of the computational problem that current tractography approaches strive to solve.

The large number of invalid bundles that were systematically identified by most state-of-the-art algorithms can only be resolved by substantial methodological innovation. Several directions of current research might improve the specificity of tractography. Streamline filtering techniques can optimize the signal prediction error in order to reduce tractography biases (14,16,56). Such techniques can increase the reproducibility and quality of tractograms. Teams 13, 16 and 17 applied such filtering techniques, showing a positive effect (albeit not significant) on the valid connection ratio, the invalid bundle count and the overreach score, at the expense of decreasing valid bundle counts and decreasing overlap. The mentioned techniques are part of the more general trend to integrate non-local as well as advanced diffusion microstructure modeling information that goes beyond the raw directional vectors (57–62). Recent advances in machine-learning-driven tractography also show high potential in improving the specificity of tractograms (63). Future versions of our phantom will be generated with multiple b-values, better signal-to-noise ratio (SNR) and fewer artifacts to further encourage research in these directions.

In addition, tractography should employ reliable anatomical priors, such as from animal experiments, for optimal guidance. While manual or automated clean-up of streamlines may help (see our results), the real challenge is our limited knowledge of the anatomy to be reconstructed. Currently, post-mortem dissection with Klingler’s method reveals the macroscopic organization of the human brain white matter (11,64–66). In the future, the community will have to gain further insights into the underlying principles of white matter organization and increasingly learn how to leverage such information for tractography (1,67,68).

Another limitation of tractography as emphasized by our results is highly relevant for the field of connectomics: The traditional metrics that require streamlines to exactly end in head or tail regions of a bundle are far too restrictive for bundle dissection and connectivity assessment. None of the submissions generated exact overlaps of streamlines with ground truth bundles and dilated endpoint masks. This finding raises an important warning for connectomics and structural connectivity studies, where a voxel-wise definition of parcellations on the T1 image is the state-of-the-art method for selecting relevant streamlines. This finding is in line with previous reports which found termination of tracts in the grey matter to be inaccurate (6). Future versions of our phantom will include ground-truth parcellations of the white matter/gray matter cortical band to encourage further developments for tackling these problems and extend the evaluation method to apply to graph theory metrics.

DWI is the only tool to map long-range structural brain connectivity *in vivo* and is essential for comparing brains, detecting differences and simulating functions (69). However, our findings should foster the development of novel tractography methods that are carefully evaluated using the presented approach. The most important goal for the next generation of tractography algorithms will no longer be to find existing valid connections, but to reconstruct the full spatial extent of tracts while controlling for the many false-positive connections polluting the tractograms. A tighter integration of advanced diffusion microstructure modeling and multi-modality imaging in tractography should help resolve ambiguities in the signal and overcome current limitations of tractography (62,70). Fundamentally, tractography will require severe methodological innovation to become tractable (1,8).

## Online Methods

### Generation of ground truth fiber bundles

The set of ground truth long-range fiber bundles was designed to cover the whole human brain and feature many of the relevant spatial configurations, such as crossing, kissing, twisting and fanning fibers, thus representing the morphology of the major known *in vivo* fiber bundles. The process to obtain these bundles consisted of three steps. First, a whole-brain global tractography was performed on a high quality *in vivo* diffusion-weighted image. Then, 25 major long-range bundles were manually extracted from the resulting tractogram. In the third step, these bundles were refined to obtain smooth and well defined bundles.

We chose one of the diffusion-weighted datasets included in the Q3 data release of the Human Connectome Project (71), subject 100307, to perform whole-brain global fiber tractography (57,72). Amongst other customizations, the HCP scanners are equipped with a set of high-end gradient coils, enabling diffusion encoding gradient strengths of 100 mT m^−1^. By comparison, most standard magnetic resonance scanners feature gradient strengths of about 30 to 40 mT m^−1^. This hardware setup allows the acquisition of datasets featuring exceptionally high resolutions (1.25 mm isotropic, 270 gradient directions) while maintaining an excellent SNR. All datasets were corrected for head motion, eddy currents and susceptibility distortions and are, in general, of very high quality (73–77). Detailed information regarding the employed imaging protocols as well as the datasets themselves may be found on http://humanconnectome.org.

Global fiber tractography was performed using *MITK Diffusion* (78) with the following parameters: 900,000,000 iterations, a particle length of 1 mm, a particle width of 0.1 mm and a particle weight of 0.002. Furthermore, we repeated the tractography six times and combined the resulting whole-brain tractograms into one large dataset consisting of over five million streamlines. The selected parameters provided for a very high sensitivity of the tractography method. The specificity of the resulting tractogram was of lesser concern since the tracts of interest were extracted manually in the second step.

Bundle segmentation was performed by an expert radiologist using manually placed inclusion and exclusion regions of interest (ROI). We followed the concepts introduced in (45) for the ROI placement and fiber extraction. 25 bundles were extracted, covering association, projection and commissural fibers across the whole brain (Figure 1): corpus callosum (CC), left and right cingulum (Cg), Fornix (Fx), anterior commissure (CA), left and right optic radiation (OR), posterior commissure (CP), left and right inferior cerebellar peduncle (ICP), middle cerebellar peduncle (MCP), left and right superior cerebellar peduncle (SCP), left and right parieto-occipital pontine tract (POPT), left and right cortico-spinal tract (CST), left and right frontopontine tracts (FPT), left and right inferior longitudinal fasciculus (ILF), left and right uncinate fasciculus (UF) and left and right superior longitudinal fasciculus (SLF). As mentioned in the Discussion section, the inferior fronto-occipital fasciculus (IFOF), the middle longitudinal fasciculus (MdLF) as well as the middle and inferior temporal projections of the arcuate fasciculus (AF) were not included.

After manual extraction, the individual long-range bundles were further refined to serve as ground truth for the image simulation as also shown in Figure 1. The original extracted tracts featured a large number of prematurely ending fibers and the individual streamlines were not smooth. To obtain smooth tracts without prematurely ending fibers, we simulated a diffusion-weighted image from each original tract individually using *Fiberfox* (www.mitk.org (35)). Since no complex fiber configurations, such as crossings, were present in the individual tract images and no artifacts were simulated, it was possible to obtain very smooth and complete tracts from these images with a simple tensor-based streamline tractography. Supplementary Figure 6 illustrates the result of this refining procedure on the left CST.

### Simulation of phantom images with brain-like geometry

The phantom diffusion-weighted images (Supplementary Video 3) were simulated using Fiberfox (www.mitk.org (35)), which is available as open-source software. We employed a four-compartment model of brain tissue (intra and inter-axonal), grey matter (GM) and cerebrospinal fluid (CSF) (35). The parameters for simulation of the four-compartment diffusion-weighted signal were chosen to obtain representative diffusion properties and image contrasts (compare (79) for details on the models). The *intra-axonal compartment* was simulated using the stick model with a T2 relaxation time of 110 ms and a diffusivity of 1.2 × 10^−9^ m^2^/s. The *inter-axonal compartment* was simulated using the zeppelin model with a T2 relaxation time of 110 ms, an axial diffusivity of 1.2 × 10^−9^ m^2^/s and a radial diffusivity of 0.3 × 10^−9^ m^2^/s. The *grey matter compartment* was simulated using the ball model with a T2 relaxation time of 80 ms and a diffusivity of 1.0 × 10^−9^ m^2^/s. The *CSF compartment* was also simulated using the ball model with a T2 relaxation time of 2500 ms and a diffusivity of 2.0 × 10^−9^ m^2^/s.

Using Fiberfox, one set of diffusion-weighted images and one T1-weighted image were simulated. The final datasets as well as all files needed to perform the simulation are available online (Supplementary Data 1).

The acquisition parameters that we report below were chosen to simulate images that are representative for a practical (e.g., clinical) setting, specifically a 5-to-10-minute single shot EPI scan with 2 mm isotropic voxels, 32 gradient directions and a b-value of 1000 s/mm^2^. The chosen acquisition setup represents a typical scenario for an applied tractography study and embodies a common denominator supported by the large majority of methods. Since acquisitions with higher b-values, more gradient directions and fewer artifacts are beneficial for tractography, we additionally report a least upper bound tractography performance under perfect image quality conditions using a dataset that directly contains ground truth fiber orientation distribution functions and no artifacts (Supplementary Figure 5 and Supplementary Notes 3).

The parameters are a matrix size of 90×108×90, echo time (TE) 108 ms, dwell time 1 ms; T2’ relaxation time 50 ms. The simulation corresponded to a single-coil acquisition with constant coil sensitivity, no partial Fourier and no parallel imaging. Phase encoding was posterior-anterior. Two unweighted images with posterior-anterior/anterior-posterior phase encoding were also generated.

Since *Fiberfox* simulates the actual k-space acquisition, it was possible to introduce a number of common artifacts into the final image. Complex *Gaussian noise* was simulated yielding a final SNR relative to the mean white matter baseline signal of about 20. 10 *spikes* were distributed randomly throughout the image volumes (Supplementary Figure 7a). *N/2 ghosts* were simulated (Supplementary Figure 7b). Distortions caused by *B_0_ field inhomogeneities* are introduced using an existing field map measured in a real acquisition and registered to the employed reference HCP dataset (Supplementary Figure 7c). *Head motion* was introduced as random rotation (+−4° around z-axis) and translation (+−2mm along x-axis) in three randomly chosen volumes. Volume 6 was rotated by 3.36° and translated by −1.74 mm, volume 12 was rotated by 1.23° and translated by −0.72 mm, and volume 24 was rotated by −3.12° and translated by −1.55 mm.

The image with the T1-like contrast was generated at an isotropic resolution of 1 mm, an SNR of about 40 and no further artifacts as an anatomical reference.

### Performance metrics and evaluation

The Tractometer definition of a valid connection is extremely restrictive for current tractography algorithms, as it requires streamlines 1) not to exit the area of the ground truth bundle at any point and 2) to terminate exactly within the endpoint region that is defined by the dilated ground truth fiber endpoints (Supplementary Figure 8 and Supplementary Figure 9) (43). Hence, we propose an alternative definition that relaxes these strict criteria based on robust shape distance measures (80) and clustering between streamlines (81), as detailed in Supplementary Notes 1. The bundle-specific thresholds were manually configured to account for bundle shape and proximity to other bundles. The following distances were used, with identical distances on both sides for lateralized bundles: 2 mm for CA and CP; 3 mm for CST and SCP; 5 mm for Cingulum; 6 mm for Fornix, ICP, OR and UF; 7 mm for FPT, ILF and POPT; 10 mm for CC, MCP and SLF. The full script used to run this bundle recognition implementation was based on the *DIPY library* (82) (www.dipy.org) and is available online (Supplementary Software 1).

Once valid connections are identified, the remaining streamlines can be classified into *invalid connections* and *non-connecting streamlines.* The details of this procedure are described in Supplementary Notes 1. We clustered the remaining invalid streamlines using a QuickBundles-based clustering algorithm (81). The best matching endpoint regions for each resulting cluster were identified by majority voting of the contained streamlines. If multiple clusters were assigned to the same pair of regions, they were merged. Streamlines that were not assigned to any cluster or that fell below a length threshold were labelled as non-connecting.

On the basis of this classification of streamlines, the following metrics were calculated:

- Valid connection ratio (VC): number of valid connections / total number of streamlines (percentage between 0 and 100).
- Valid bundles (VB): For each bundle that has a valid streamline associated with it, this counter is incremented by one (integer number between 0 and 25).
- Invalid bundles (IB): With 25 bundles in the ground truth, each having two endpoint regions, there are 1,275 possible combinations of endpoint regions. Taking the 25 valid bundles out of the equation, 1,250 potential invalid bundles remain (integer number between 0 and 1,250).
- Overlap: Proportion of the voxels within the volume of a ground truth bundle that is traversed by at least one valid streamline associated with the bundle. This value shows how well the tractography result recovers the original volume of the bundle (percentage between 0 and 100).
- Overreach: Fraction of voxels outside the volume of a ground truth bundle that is traversed by at least one valid streamline associated with the bundle over the total number of voxels within the ground truth bundle. This value shows how much the valid connections extend beyond the ground truth bundle volume (percentage between 0 and 100). This value is always zero for the traditional definition of a valid connection but can be non-zero for the relaxed evaluation.

### Statistical multi-variable analysis

Effects of the experimental settings were investigated in a multivariable linear mixed model. The experimental variables describing the methods used for the different submissions were included as fixed effects (Figure 2b). The valid connection ratio, the valid bundle count, the invalid bundle count, the bundle overlap percentage and the bundle overreach percentage were modeled as dependent variables, each of which is used for the calculation of a separate model. The submitting group was modeled as a random effect. The software *SAS 9.2*, Proc Mixed, SAS Institute Inc., Cary, NC, USA was used for the analysis.

## Acknowledgements

This work was supported by the German Research Foundation (DFG), grants MA 6340/10-1, MA 6340/12-1 and the NSERC Discovery Grant program as well as the institutional Université de Sherbrooke Research Chair in Neuroinformatics. C.M.W.T. is supported by a grant (No. 612.001.104) from the Physical Sciences division of the Netherlands Organization for Scientific Research (NWO). The research of H.Y.M., S.D., S.S., A.M.H. and A.L. is supported by VIDI Grant 639.072.411 from NWO. The research of F.G. was funded by the Chinese Scholarship Council (CSC). M.C. is supported by the Alexander Graham Bell Canada Graduate Scholarships-Doctoral Program (CGS-D3) from the Natural Sciences and Engineering Research Council of Canada (NSERC). C.C.H. is supported by DFG SFB grants 936/A1, Z3 and TRR 169/A2.

## Author contributions

K.M.H., M.D. and J-C.H. performed the data analysis and wrote the paper with input from all authors. P.N. and B.S. designed the phantom. P.N. and J-C.H. supported the data analysis and J-C.H handled the Tractometer scoring and evaluation metrics proposed. M-A.C and E.G. developed the clustering and bundle recognition algorithm for the relaxed scoring system. K.M.H., P.N., J-C.H., E.C., A.D., T.D., B.S. and M.D. coordinated the tractography challenge at the International Society for Magnetic Resonance in Medicine (ISMRM) 2015 Diffusion Study Group meeting. T.H-L. set up the multivariable statistical model. P.N. wrote parts of the Online Methods. L.P. and C.C.H. were mentors in the discussion of the paper and neuroanatomical as well as neuroscientific context. Submissions were made by the following teams: J.Z. team 1; M.C. and C.M.W.T. team 2; F-C.Y. team 3; Y-C.L. team 4; Q.J. team 5; D.Q.C. team 6; Y.F., C.G., Y.W., J.M., H.R., Q.L. and C-F.W. team 7; S.D-G., J.O.O.G., M.P., S.S-J. and G.G. team 8; S.S-J., F.R. and J.S. team 9; C.M.W.T., F.G., H.Y.M., S.D., M.F., A.M.H. and A.L. team 10; S.S-J., G.G. and F.R. team 11; J.O.O.G., M.P., G.G. and F.R. team 12; A.B., B.P., C.B., M.D., S.B. and J.D. team 13; A.S., R.V., A.C., A.Q. and J.Y. team 14; A.R.K., W.H. and S.A. team 15; D.R., M.B., A.A., O.E., A.L. and J-P.T. team 16; D.R., M.B., A.A., O.E., A.L. and J-P.T. team 17; H.E.C., B.L.O., B.M. and M.S.N. team 18; F.P., G.P., J.E.V-R., J.G. and P.M.T. team 19; F.D.S.R., P.L.L., L.M.L., R.B. and F.D’A team 20.

## Additional information

Supplementary information is available in the online version of the paper. Correspondence and requests for materials should be addressed to K.M.H. or M.D.

## Competing financial interests

The authors declare no competing financial interests.

